# IMPDH inhibitors for anti-tumor therapy in tuberous sclerosis complex

**DOI:** 10.1101/835199

**Authors:** Alexander J. Valvezan, Spencer K. Miller, Molly C. McNamara, Margaret E. Torrence, John M. Asara, Elizabeth P. Henske, Brendan D. Manning

## Abstract

**Purpose:** mTORC1 is a master regulator of anabolic cell growth and proliferation that is activated in the majority of human tumors. We recently demonstrated that elevated mTORC1 activity in cells and tumors can confer dependence on IMPDH, the rate-limiting enzyme in *de novo* guanylate nucleotide synthesis, to support increased ribosome biogenesis and cell viability. Pharmacological agents that inhibit IMPDH, such as mizoribine and mycophenolic acid (CellCept), are in wide clinical use as immunosuppressants. However, whether these agents can be repurposed for anti-tumor therapy requires further investigation in preclinical models, including direct comparisons to identify the best candidate(s) for advancement.

**Experimental Design:** Distinct IMPDH inhibitors were tested on cell and mouse tumor models of tuberous sclerosis complex (TSC), a genetic tumor syndrome featuring widespread lesions with uncontrolled mTORC1 activity. Growth and viability were assessed in cells and tumors lacking the TSC2 tumor suppressor, together with drug pharmacokinetics and pharmacodynamics, target inhibition, and effects on tumor, tissue, and plasma metabolic biomarkers.

**Results:** Mizoribine, used throughout Asia, exhibited greater selectivity in specifically targeting TSC2-deficient cells with active mTORC1 compared to the FDA-approved IMPDH inhibitors mycophenolic acid or ribavirin, or approved inhibitors of other nucleotide synthesis enzymes. In distinct tumor models, mizoribine demonstrated robust anti-tumor efficacy that is superior to mycophenolic acid, despite similar immunosuppressive effects.

**Conclusions:** These results provide pre-clinical rationale for repurposing mizoribine as an anti-tumor agent in tumors with active mTORC1, such as in TSC. Our findings also suggest that IMPDH inhibitors should be revisited in cancer models where MMF has shown modest efficacy.

**Statement of translational relevance:** IMPDH inhibitors have been used clinically for decades as safe and effective immunosuppressants. Recent studies in pre-clinical tumor models establish IMPDH as a viable target for anti-tumor therapy, but the relative efficacies of approved IMPDH inhibitors in tumors have not been directly compared. Our data demonstrate a clear advantage of the IMPDH inhibitor mizoribine, used clinically throughout Asia, over the FDA-approved IMPDH inhibitor mycophenolate mofetil (or CellCept, a prodrug of mycophenolic acid) in mouse models of tuberous sclerosis complex (TSC) exhibiting mTORC1-driven tumor growth. While these IMPDH inhibitors elicit similar immunosuppressive effects, mizoribine has far superior anti-tumor activity in these models, indicating the potential for repurposing this drug for TSC and perhaps cancer treatment. We also identify the purine synthesis intermediate AICAR as an *in vivo* metabolic biomarker specific for effective inhibition of IMPDH with mizoribine, which can be readily detected in blood plasma shortly after mizoribine administration.

## Introduction

Tumor cells continuously synthesize new biomass to support their uncontrolled proliferation. This is often achieved, in part, through activation of mechanistic Target of Rapamycin Complex 1 (mTORC1), a central regulator of anabolic cell growth that is active in the majority of human cancers, across nearly all lineages^1, 2^. mTORC1 induces a coordinated anabolic program to stimulate synthesis of the major macromolecules required for cell growth, including proteins, lipids, and nucleic acids^3^. mTORC1 also stimulates flux through metabolic pathways that support macromolecule synthesis, including *de novo* purine and pyrimidine nucleotide synthesis pathways, glycolysis, and the oxidative branch of the pentose phosphate pathway^4–7^. Given its role in tumor cell growth and metabolism, there has been much interest in targeting mTORC1 in cancer. However, with over 1000 clinical trials to date, FDA-approved mTORC1 inhibitors (rapamycin/sirolimus and its analogs) have shown limited efficacy as single agent therapies, consistent with their established growth-slowing, cytostatic effects rather than the cytotoxic effects needed to eliminate tumor cells^8^.

Rapamycin and its analogs are currently in frequent use for treatment of tumors in tuberous sclerosis complex (TSC), a genetic syndrome caused by loss of function mutations in the *TSC1* or *TSC2* tumor suppressor genes, which encode the essential components of the TSC protein complex (TSC complex)^9^. The TSC complex inhibits the Ras-related GTPase Rheb, which is an essential upstream activator of mTORC1, thus tumors in TSC patients are driven by robust, uncontrolled mTORC1 activation^10^. TSC is a pleiotropic disorder in which patients commonly develop neurological phenotypes such as epilepsy, autism, and a variety of cognitive and behavioral manifestations (collectively referred to as TSC-associated neuropsychiatric disorders or TAND), accompanied by widespread tumor development across multiple organ systems including, but not limited to, the brain (subependymal giant cell astrocytomas), heart (rhabdomyomas), kidney (angiomyolipomas), skin (fibromas), and lung (lymphangioleiomyomatosis (LAM))^11^. LAM is a proliferative and destructive lung disorder that can lead to respiratory failure, is nearly exclusive to women, and arises in both TSC patients and sporadically through inactivating mutations in *TSC1* or *TSC2*^12^. Rapamycin and its analogs can slow or shrink tumors in TSC but do not kill tumor cells. Thus, tumors are not eliminated by these agents and regrow rapidly when treatment is discontinued^13, 14^. Mutations in *TSC1* and *TSC2* are also found in sporadic cancers, with the highest frequency being in bladder cancer and hepatocellular carcinoma^15, 16^. Thus, there is an unmet clinical need to selectively induce cell death in TSC1/2-deficient tumors. Finally, it is worth noting that the primary route to uncontrolled mTORC1 activity in human cancers is through aberrant inhibition of the TSC complex, as some of the most commonly altered oncogenes (e.g., *PIK3CA*, *KRAS*) and tumor suppressors (*PTEN, NF1, LKB1*) lie within signaling pathways that converge on regulation of the TSC complex to control mTORC1^17^.

As an alternative to inhibiting mTORC1 itself in tumors, it might be possible to elicit sustained anti-tumor responses by targeting specific processes within the downstream metabolic network activated as part of the cell growth program of mTORC1. Indeed, we previously demonstrated that aberrant mTORC1 activation increases dependence on nucleotide synthesis for cell viability. In cells and tumors with active mTORC1, nucleotide synthesis pathways are required to meet the increased demand for nucleotides that comes from mTORC1 stimulating ribosomal RNA synthesis^18^, exemplifying the importance of mTORC1 coupling distinct metabolic pathways within the broader anabolic program it induces. This metabolic vulnerability can be exploited by inhibiting inosine monophosphate dehydrogenase (IMPDH), the rate-limiting enzyme in *de novo* guanylate nucleotide synthesis. IMPDH inhibition was found to cause DNA replication stress preferentially in cells and tumors with active mTORC1, which culminates in DNA damage and apoptosis^18^. Thus, in contrast to mTORC1 inhibitors, IMPDH inhibition is selectively cytotoxic to cells with uncontrolled mTORC1 signaling, and co-treatment with rapamycin actually counteracts these effects of IMPDH inhibitors. IMPDH2 is frequently upregulated in human cancers, including a subset of small cell lung cancers where IMPDH inhibition was recently shown to have anti-tumor efficacy in preclinical tumor models^19^. IMPDH inhibition has also shown efficacy in xenograft tumors established using human glioblastoma, leukemia, lymphoma, pancreatic adenocarcinoma, and colon adenocarcinoma cell lines^20–22^. If these preclinical findings are to be translated into clinical anti-tumor therapy, it is essential to directly compare the efficacy of available IMPDH inhibitors.

Activated lymphocytes require mTORC1 signaling and *de novo* nucleotide synthesis pathways for their rapid expansion and thus inhibitors of these pathways, including rapamycin and IMPDH inhibitors, are used clinically as immunosuppressants. IMPDH inhibitors are well-tolerated, especially when compared to classical chemotherapeutic agents that directly target dNTP or DNA synthesis^23^, making them attractive for potential repurposing. Of the clinically approved IMPDH inhibitors: mizoribine (Bredinin) is a natural purine analog used predominantly in Asia for autoimmune disorders and preventing organ rejection after transplant, mycophenolic acid (MPA, Myfortic) and its orally bioavailable prodrug mycophenolate mofetil (MMF, Cellcept) are FDA approved and structurally distinct from mizoribine but used for similar indications as mizoribine, and ribavirin (Rebetol, Copegus) is an FDA approved purine analog that inhibits IMPDH, but unlike mizoribine, ribavirin incorporates into RNA, and is used as an anti-viral agent^24, 25^.

We hypothesized that these IMPDH inhibitors may not be equivalent in their ability to selectively target cells with active mTORC1 and prevent mTORC1-driven tumor growth at clinically relevant doses. This hypothesis was tested in cells and tumors lacking a functional TSC complex, which provide well-established models of mTORC1-driven growth, as well as translationally relevant disease models of TSC.

## Materials and Methods

### Cell Lines and culture conditions

*Tsc2^+/+^; Trp53^−/−^ and Tsc2^−/−^; Trp53^−/−^* MEFs^26^, *Tsc2^−/−^* 105Ks stably expressing empty vector or TSC2^27^, 621-101 cells stably expressing empty vector or TSC2^28^, HeLa cells stably expressing shRNAs targeting luciferase or TSC2^29^, and *Tsc2^−/−^*ELT3 cells^30^ were described previously. Cells were grown in DMEM (VWR #45000-312) plus 10% heat-inactivated fetal bovine serum (ThermoFisher Scientific #10437-028) and 1% penicillin/streptomycin (Corning #30-002-Cl).

### Chemical Compounds

Where indicated, the following compounds were added into the cell culture medium at final concentrations stated in the figures and legends: mizoribine (Selleckchem #S1384 and Sigma #M3047), mycophenolic acid (Sigma #M3536), mycophenolate mofetil (Selleckchem #S1501), ribavirin (Sigma #R9644), AVN-944 (Chemietek #CT-AVN944), rapamycin (EMD Millipore #53123-88-9), methotrexate (Sigma #A6770), 6-mercaptopurine (Sigma #852678), A771726 (Calbiochem #100128), Brequinar (Sigma #SML0113), pyrazofurin (Sigma #SML1502), 3-deazauridine (Santa Cruz #sc-394445), 5-fluorouracil (Tocris #3257), guanosine (Sigma #G6752).

### siRNA Transfections

Control non-targeting siRNA (GE Dharmacon D-001810-10-50) or siRNA targeting ADK (GE Dharmacon L-062728-00-0005) were transfected at 12.5 nM using Lipofectamine RNAiMAX (ThermoFisher Scientific #13778150) according to manufacturer’s instructions.

### Cell Viability Assays

Cell viability was determined by trypan blue (Sigma #T8154) exclusion using a hemocytometer. Where indicated, viable cell counts were measured using the Cell Titer Glo Luminescent Cell Viability Assay (Promega #G7573) according to manufacturer’s instructions. Annexin V/PI staining was performed using the Dead Cell Apoptosis Kit (ThermoFisher Scientific #V13245), according to manufacturer’s instructions. Staining was measured with a Becton Dickinson LSR Fortessa flow cytometer, and analyzed with FlowJo Version 10.2 software.

### Immunoblotting and Antibodies

Cells were lysed in buffer containing 20 mM Tris pH 7.5, 140 mM NaCl, 1 mM EDTA, 10% glycerol, 1% Triton X-100, 1 mM DTT, 50 mM NaF, protease inhibitor cocktail (Sigma #P8340) and phosphatase inhibitor cocktail #2 (Sigma #P5726) and #3 (Sigma #P0044) used at 1:100. Western blots were performed using the following antibodies at 1:1000 dilution unless otherwise indicated: phospho-p70 S6 Kinase T389 (CST #9234), p70 S6 Kinase (CST #2708), TSC2 (CST #4308 and #3612), Rheb (CST# 13879), GAPDH (CST #5174), Cleaved Caspase-3 D175 (CST #9664), ADK (Santa Cruz #sc-514588, 1:500), phospho-Chk1 S345 (CST #2348), Chk1 (CST #2360), phospho-Histone H2A.X S139 (CST #9718), and Histone H2A.X (CST #7631).

### Mouse Studies

All animal procedures were approved by the Harvard Institutional Animal Care and Use Committee. For xenograft tumor studies, 2.5 million 105K cells or ELT3 cells expressing luciferase^31^ were subcutaneously injected in a 1:1 mixture with matrigel (BD #356237) into the flank of 6-7 week old NOD.Cg-Prkdc^scid^ Il2rg^tm1Wjl^/SzJ (NSG) mice (Jackson Laboratory #005557) or C57BL/6J mice (Jackson Laboratory #000664). Treatment with mizoribine (Selleckchem #S1384), rapamycin (LC laboratories #R-5000), or mycophenolate mofetil (Sigma #1448956) began 4 weeks later, when tumors first became palpable, using doses indicated in the figures and legends. Mice were assigned to treatment groups based on their tumor size, so that the average tumor size per group was the same among all groups at the start of treatment. Tumor size was measured every three days using digital calipers. *Tsc2^+/-^* mice on the A/J strain background were described previously^32, 33^ and were treated through the TS Alliance Preclinical Consortium at PsychoGenics. Mice were randomly assigned for treatment with vehicle or MMF (75 mg/kg/day) by oral gavage for 1 month beginning at 7 months of age. Kidneys were resected, fixed in 4% paraformaldehyde for 24 hrs and embedded in paraffin. Kidneys were sectioned and tumor volume quantified as previously described^32^.

### Immunohistochemistry

Immunohistochemistry (IHC) staining was performed on formalin-fixed, paraffin-embedded tissue sections using a previously described staining protocol^18^ with the following antibodies: phospho-S6 S235/236 (CST #4858 1:400), ADK (Abcam #ab204430 1:20), Ki67 (Abcam #ab16667 1:100), Cleaved Caspase-3 (CST #9664 1:100). Human pulmonary lymphangioleiomyomatosis and renal angiomyolipoma patient samples were collected according to IRB protocol approved by Brigham and Women’s Hospital.

### Plasma collection from mice and blood cell counts

Blood was drawn by retro-orbital insertion of a heparinized microcapillary tube (Fisher #22-362-566), collected in EDTA-coated microtainer tubes (BD #365973), centrifuged at 3000xg for 10 min at room temperature, and plasma was removed. Blood cell counts were determined using a Mascot HemaVet 950FS Hematology Analyzer.

### Metabolite Analyses by LC-MS/MS

Metabolites were extracted using 80% methanol and dried under nitrogen gas for targeted tandem mass spectrometry (LC-MS/MS) profiling via selected reaction monitoring (SRM) with polarity switching on a 6500 QTRAP mass spectrometer (AB/SCIEX) as previously described^5, 34, 35^. For LC-MS/MS analysis, metabolites were extracted from either 80 µl plasma, 0.1 - 0.5 g tissue, or one 10 cm cell culture dish. Mizoribine was measured in negative ion mode using Q1/Q3 SRM transitions of 258.2/126 and mycophenolic acid in positive ion mode using Q1/Q3 SRM transitions of 321.1/206.9. Data were analyzed by calculating the Q3 peak areas using MultiQuant 3.0 software (AB/SCIEX).

### Quantification and statistical analysis

Graphical data are presented as mean ± SEM. p values for pairwise comparisons were determined using an unpaired two-tailed Student’s t test. Statistical details for individual experiments can be found in their respective figure legends.

## Results

### Mizoribine is the most selective nucleotide synthesis inhibitor for targeting TSC2-deficient cells *in vitro*

We previously demonstrated that uncontrolled mTORC1 activation stemming from loss of the TSC complex increases cellular dependence on nucleotide synthesis pathways for growth and viability, and that inhibitors of IMPDH induce apoptosis selectively in cells with active mTORC1^18^. To determine whether other enzymatic targets in nucleotide synthesis pathways can elicit a similar preferential response, littermate-derived *Tsc2^+/+^* and *Tsc2^−/−^* MEFs were treated with a panel of available inhibitors of enzymes in the *de novo* purine and pyrimidine synthesis and salvage pathways (Supplementary Fig. S1A-S1G). However, none preferentially inhibited the proliferation of *Tsc2^−/−^*cells compared to *Tsc2^+/+^*, including inhibitors of dihydrofolate reductase (methotrexate), hypoxanthine-guanine phosphoribosyltransferase (6-mercaptopurine), dihydroorotate dehydrogenase (A771726 and brequinar), uridine monophosphate synthase (pyrazofurin), cytidine triphosphate synthase (3-deazauridine), and thymidylate synthase (5-fluorouracil). Thus, we focused on structurally distinct inhibitors of IMPDH to identify the compound that most selectively inhibits the viability of cells with uncontrolled mTORC1 signaling.

The clinically used compounds mizoribine, MPA, and ribavirin, which inhibit both IMPDH isoforms (IMPDH1 and IMPDH2), all preferentially reduced the proliferation of *Tsc2^−/−^* cells compared to *Tsc2^+/+^*, with mizoribine exhibiting the most selective effects on *Tsc2^−/−^* cells and ribavirin the least (Fig. 1A). At effective doses, MPA and ribavirin were more generally cytotoxic to both wild-type and Tsc2^−/−^ cells relative to mizoribine. A fourth IMPDH inhibitor that is not in clinical use, AVN-944^36^, did not exhibit preferential inhibition of *Tsc2^−/−^*cell growth (Supplementary Fig. S1H). Mizoribine also exhibited greater selectivity compared to MPA in three isogenic pairs of TSC2-deficient or expressing cell lines: a murine *Tsc2^−/−^* renal tumor-derived cell line (105K) and a human *TSC2^−/−^* renal angiomyolipoma-derived cell line (621-101), both stably reconstituted with either wild type TSC2 or empty vector, and HeLa cells with stable shRNA-mediated knockdown of TSC2 or non-targeting control (Fig. 1B, Supplementary Fig. S1I,J). Importantly, these effects on viable cell number reflect selective induction of apoptosis by mizoribine in *Tsc2^−/−^*cells, as measured by caspase-3 cleavage and annexin V/propidium iodide staining (Figure 1C,D). Consistent with previous reports^37, 38^, higher doses of MPA and AVN-944 reduced mTORC1 signaling in wild-type cells, as measured by phosphorylation of the mTORC1 substrate S6 kinase (S6K), likely due to their reported effects on the protein levels of Rheb^36^, whereas mizoribine did not affect Rheb levels or mTORC1 activity (Fig. 1C).

**Figure 1.**
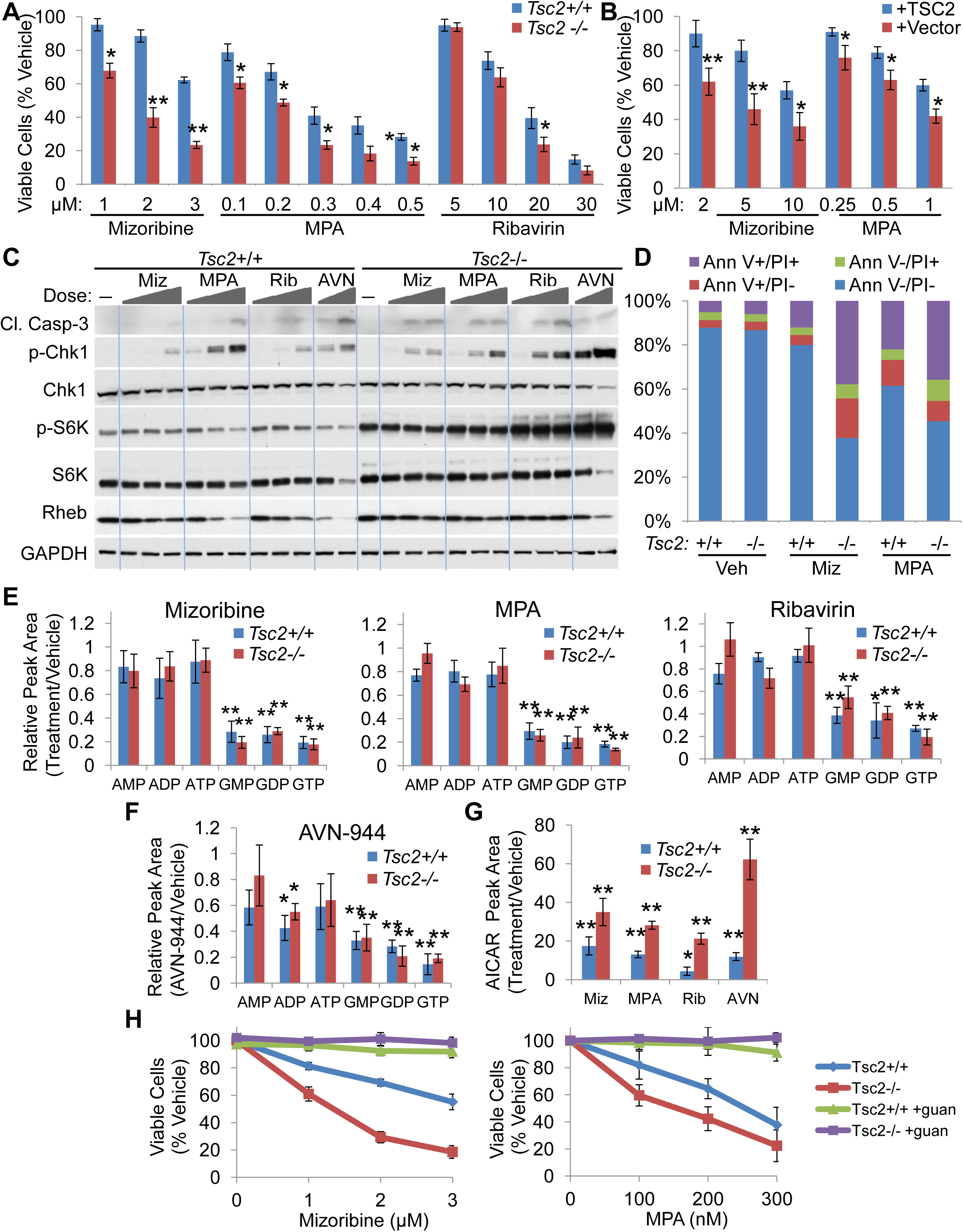
Mizoribine is the most selective IMPDH inhibitor for reducing the growth and viability of TSC2-deficient cells. (A) Littermate derived *Tsc2^+/+^* and *Tsc2*^−/−^ MEFs or (B) *Tsc2^−/−^* 105K renal tumor-derived cells stably reconstituted with empty vector or wild-type TSC2 were treated with indicated concentrations of vehicle or indicated IMPDH inhibitors for (A) 72 hrs or (B) 48 hrs. Viable cells were counted by trypan blue exclusion and graphed as percent of vehicle treated cells. (C) Cells in (A) were treated for 24 hrs with vehicle, mizoribine (Miz: 1, 2, or 3 μM), mycophenolic acid (MPA: 125, 250 or 500 nM), ribavirin (Rib: 10, 20 or 30 μM) or AVN-944 (AVN: 100 or 250 nM) followed by immunoblotting for indicated proteins. (D) Annexin V (Ann V)/Propidium Iodide (PI) staining on cells in (A) treated for 72 with vehicle, 3 μM mizoribine or 250 nM MPA. (E,F) Guanylate and adenylate nucleotides measured by LC-MS/MS in cells in (A) treated for 12 hrs with vehicle, mizoribine (2 μM), MPA (250 nM), ribavirin (20 μM) or AVN-944 (250nM) and graphed relative to vehicle. (G) Relative AICAR levels in samples from E,F. (H) Viable cell counts in cells treated for 72 with indicated concentrations of mizoribine or MPA with or without addition of exogenous guanosine (guan, 50 μM). Graphical data are presented as mean ± SEM of 3 biological replicates and are representatitve of two or more independent experiments. *p < 0.05, **p < 0.01 by two-tailed Student’s t test.

Differences in selectivity among the clinically used IMPDH inhibitors in these cell culture studies could not be explained by varying effects on purine nucleotide levels, as mizoribine, MPA, and ribavirin caused similar reduction in guanylates, with only minor effect on adenylates, at doses that yield comparable effects on *Tsc2^−/−^* cell proliferation (Fig. 1E). AVN-944 reduced guanylates but also modestly reduced adenylates (Fig. 1F). All four IMPDH inhibitors caused accumulation of aminoimidazole carboxamide ribonucleotide (AICAR), an intermediate in the *de novo* purine synthesis pathway upstream of IMPDH (Fig. 1G). Consistent with mTORC1 activation resulting in increased metabolic flux through the *de novo* purine synthesis pathway^4^, AICAR accumulated to a greater extent in *Tsc2^−/−^* cells compared to *Tsc2^+/+^*cells with all four treatments. Importantly, the effects of mizoribine and MPA on cell proliferation and apoptosis were rescued by addition of excess exogenous guanosine, which is converted to GMP independent of IMPDH, confirming that these effects on viability result from guanylate nucleotide depletion upon IMPDH inhibition (Fig. 1H, Supplementary Fig. S1K).

### Mizoribine is superior to mycophenolate in mouse TSC tumor models

Given that mizoribine and MPA were the two most selective nucleotide synthesis inhibitors for targeting TSC2-deficient cells compared to wild-type cells in culture and that they are used clinically for similar indications^39, 40^, differing from ribavirin (which has other known mechanisms of action besides IMPDH inhibition^24, 25^), we directly compared their in vivo effects in distinct TSC2-deficient xenograft models of mTORC1-driven tumor growth. Mice bearing xenograft tumors established with the *Tsc2^−/−^*renal tumor-derived 105K cell line were treated daily with vehicle, mizoribine (75 mg/kg by intraperitoneal (i.p.) injection), or the FDA approved MPA pro-drug MMF (100 mg/kg by oral gavage) for 20 days. These treatments were also compared to the mTORC1 inhibitor rapamycin (1 mg/kg MWF by i.p. injection), which has been shown previously to be effective in this tumor model and is used to treat tumors in TSC patients^41^. Treatment was initiated when tumors first became palpable and measurable (Fig. 2A). Tumor growth was completely halted by mizoribine or rapamycin treatment, while only modestly slowed by MMF (Fig. 2B). Total body weight was not affected in any treatment group (Supplementary Fig. S2A). Plasma concentrations of mizoribine and MPA were measured at the end of the treatment phase by liquid chromatography tandem mass spectrometry (LC-MS/MS) and quantified using standard curves (Supplementary Fig. S2B,C). Plasma concentrations of both compounds were well within the therapeutically achievable ranges typically used for immunosuppression in humans (Fig. 2C)^42–45^. Tumor immunohistochemical (IHC) staining revealed a slight reduction in the proliferation marker Ki67 in MMF-treated tumors compared to vehicle, but no increase in tumor cell death as determined by H&E and cleaved caspase-3 staining (Fig. 2D). Similar to cells in culture (Fig. 1C), MMF slightly reduced phosphorylation of the direct mTORC1 substrate S6K and the S6K substrate ribosomal protein S6 (Fig. 2D,E). As rapamycin treated tumors are known to resume growth when treatment is discontinued, we compared the rate of tumor regrowth in the mizoribine and rapamycin groups to assess the durability of the anti-tumor response to these compounds. Rather than resecting tumors after the final treatment, tumors in these groups were allowed to regrow for up to 38 days, until they neared maximum allowable size (Fig. 2A). Importantly, mizoribine-treated tumors regrew more slowly than those treated with rapamycin (Fig. 2B). At the end of the regrowth phase, despite not being treated for several weeks, tumors in the mizoribine group contained large necrotic and apoptotic regions, which were not observed in those initially treated with rapamycin (Fig. 2F). These data indicate that mizoribine exerts a more durable cytotoxic effect on the tumor cells in this model, as compared to the established cytostatic effects of rapamycin^8, 13, 41^.

**Figure 2.**
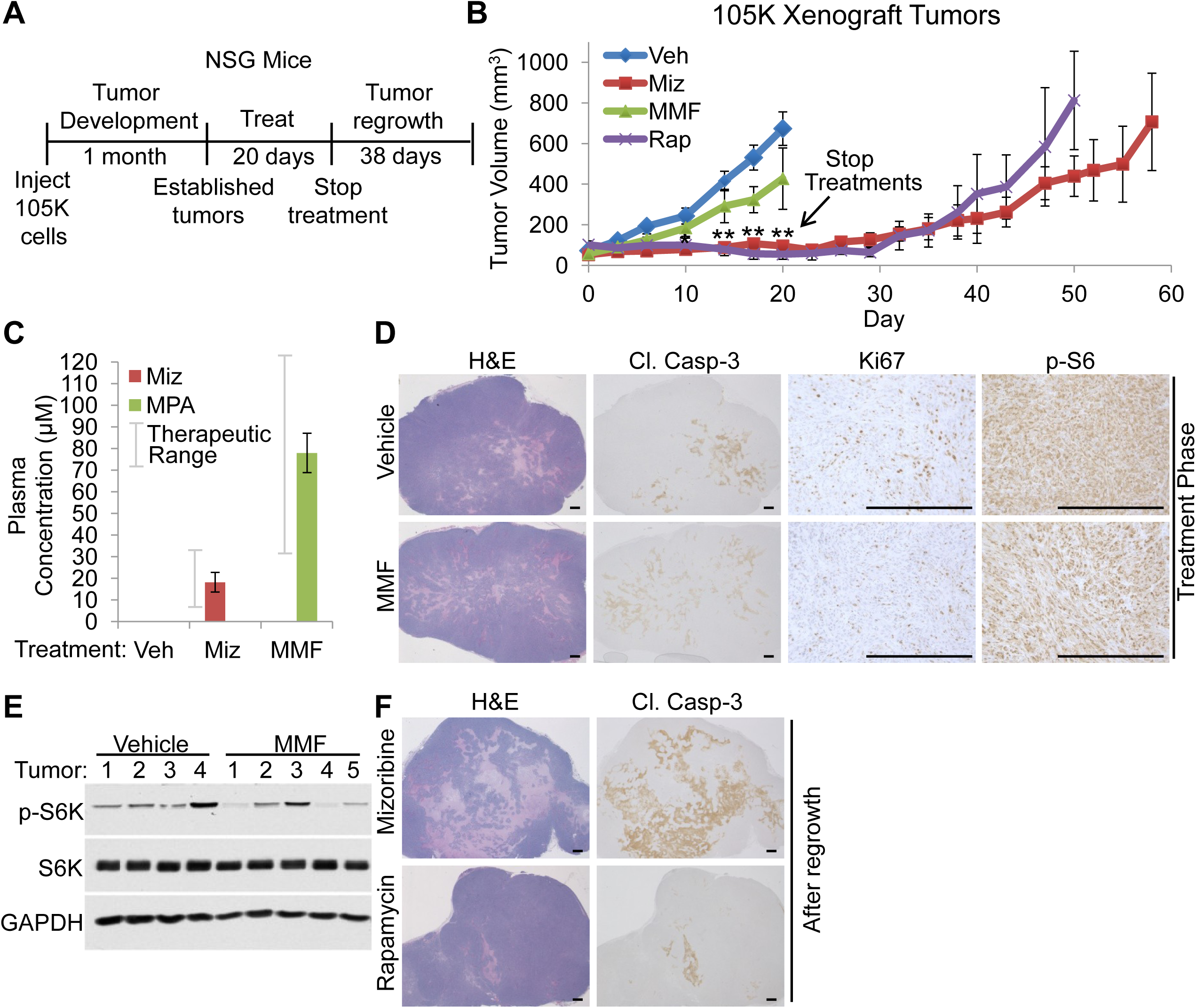
Mizoribine is superior to mycophenolate in TSC2-deficient 105K xenograft tumors at therapeutically relevant concentrations. (A) Experimental design used in (B-F). NSG mice bearing 105K cell xenograft tumors were treated for 20 days with vehicle, mizoribine (75 mg/kg/day by i.p. injection), mycophenolate mofetil (MMF, 100 mg/kg/day by oral gavage) or rapamycin (1 mg/kg Mon/Wed/Fri by i.p. injection). Treatments were discontinued after 20 days and tumors in the mizoribine and rapamycin groups were allowed to regrow. (B) Tumor volume measured every 3^rd^ day. n = 6 mice per group. (C) Mizoribine and MPA concentrations in blood plasma collected 2.5 hrs after the final treatment, measured by LC-MS/MS. n = 6 mice per group. (D) Representative H&E and immunohistochemistry staining and (E) immunoblotting on tumors from vehicle or MMF treated mice resected 3 hrs after the final treatment. (F) Representative H&E and immunohistochemistry staining on tumors from mizoribine or rapamycin treated mice resected after regrowth. Graphical data are presented as mean ± SEM of 6 mice per group. *p < 0.05, **p < 0.01 by two-tailed Student’s t test. Scale bars = 0.5 mm

Since rapamycin and its analogs are frequently prescribed to treat tumors in TSC patients, we asked whether previous rapamycin treatment would affect the response of TSC2-deficient tumors to mizoribine and MMF. Leveraging a second widely used TSC tumor model that exhibits rapid tumor regrowth upon halting rapamycin treatment, xenograft tumors were established using the TSC2-deficient rat uterine leiomyoma-derived ELT3 cell line. Mice were treated as above for 26 days with vehicle, mizoribine, or MMF, or 23 days with rapamycin, beginning when tumors first became palpable. On day 24, the rapamycin-treated mice were switched to vehicle, mizoribine, or MMF treatment for another 17 days (Fig. 3A). Consistent with findings from the 105K xenograft tumor model (Fig. 2B), mizoribine had a greater effect on ELT3 xenograft tumor growth compared to MMF, and prior rapamycin treatment did not influence this response (Fig. 3B,C). Again, these treatments did not affect total body weight (Supplementary Fig. S2D), and final plasma mizoribine and MPA concentrations were similar to those measured in the 105K tumor-bearing mice (Fig. 3D). These results indicate that the anti-tumor activity of IMPDH inhibitors, and superiority of mizoribine over MMF, are not affected by previous rapamycin treatment of TSC tumors.

**Figure 3.**
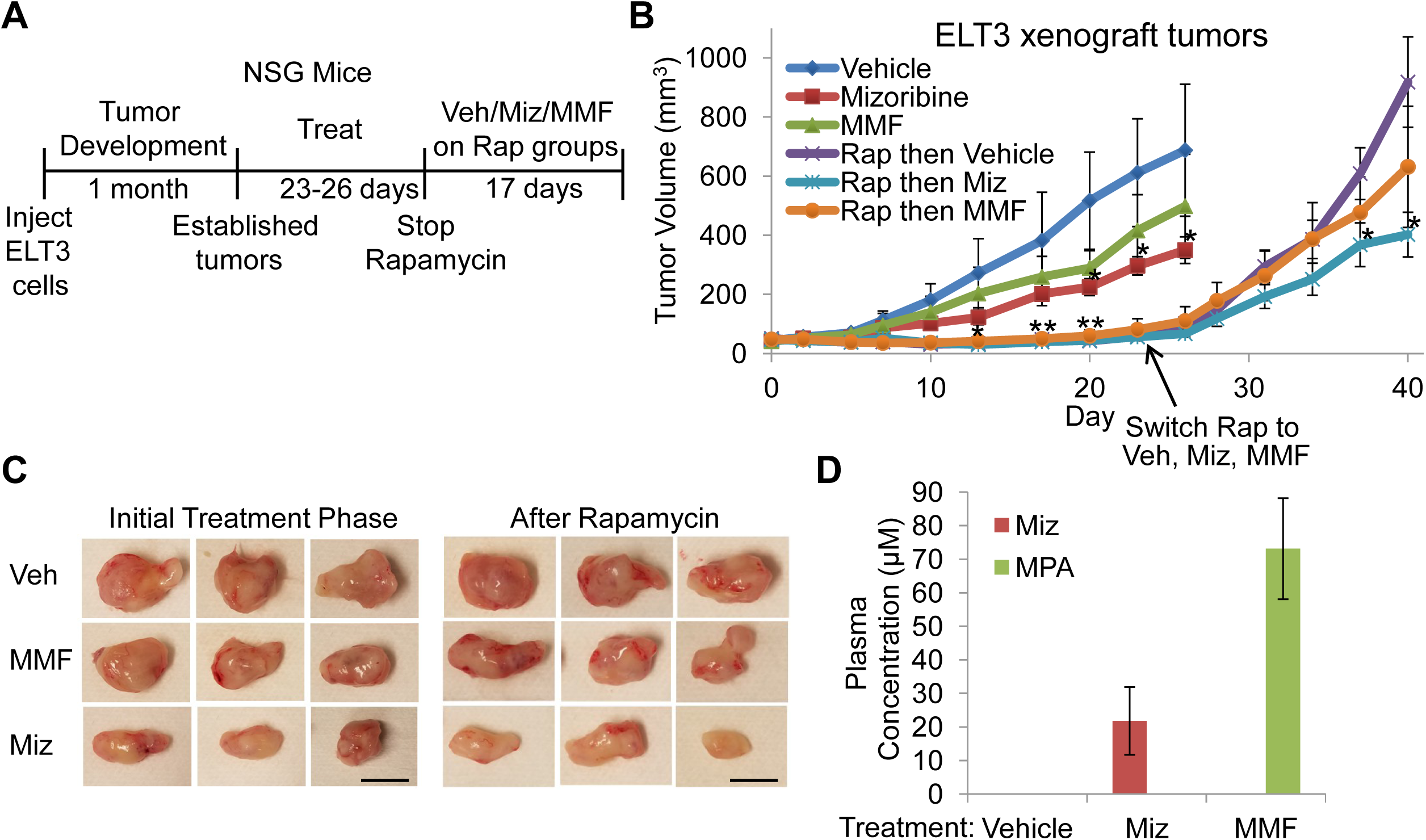
Mizoribine is superior to mycophenolate in TSC2-deficient ELT3 xenograft tumors with or without prior treatment with rapamycin. (A) Experimental design used in (B-D). NSG mice bearing ELT3 cell xenograft tumors were treated for 26 days with vehicle, mizoribine (75 mg/kg/day by i.p. injection), MMF (100 mg/kg/day by oral gavage), or 23 days with rapamycin (1 mg/kg Mon/Wed/Fri by i.p. injection). On Day 23, rapamycin treatment was discontinued and those mice were switched to vehicle, mizoribine (75 mg/kg/day by i.p. injection), or MMF (100 mg/kg/day by oral gavage) treatment for 17 days. (B) Tumor volume measured every 3^rd^ day. n = 6 mice per group. (C) Images of representative tumors from taken immediately after resection. Scale bars = 1 cm. (D) Mizoribine and MPA concentrations in blood plasma collected on Day 26, 2.5 hrs after the final treatment and measured by LC-MS/MS. n = 6 mice per group Graphical data are presented as mean ± SEM of 6 mice per group. *p < 0.05, **p < 0.01 by two-tailed Student’s t test.

### Mizoribine and mycophenolate have differential effects on metabolism *in vivo*

In attempt to determine the underlying cause of the differential anti-tumor response between mizoribine and MMF measured in these distinct TSC tumor models, we profiled steady state metabolite levels in the ELT3 xenograft tumors after 26 days of treatment using LC-MS/MS-based metabolomics (tumor metabolites significantly changed relative to vehicle shown in Fig. 4A,B). Of the tumor metabolites significantly changed in response to mizoribine and MMF, 51% and 41%, respectively, were shared between the two treatments (18 total, Fig. 4A-C). For instance, both mizoribine and MMF increased tumor putrescine and N-acetyl spermidine levels, consistent with a recent report demonstrating that the IMPDH inhibitor ribavirin can stimulate an increase in these polyamines^46^. While both drugs decreased tumor GMP levels, mizoribine had markedly stronger effects on tumor GDP and GTP than MMF, with the tumors from MMF-treated mice exhibiting modestly lower levels of adenylates relative to vehicle or mizorbine (Fig. 4D). With a 20-fold increase, AICAR was the most significantly elevated metabolite (p<0.0003) in mizoribine-treated tumors, and consistent with our previous findings, mizoribine treatment also increased plasma AICAR levels (Fig. 4E). However, in contrast to MPA-treated cells in culture (Fig. 1G), MMF did not cause significant AICAR accumulation in tumors or plasma (Fig. 4B,E).

**Figure 4.**
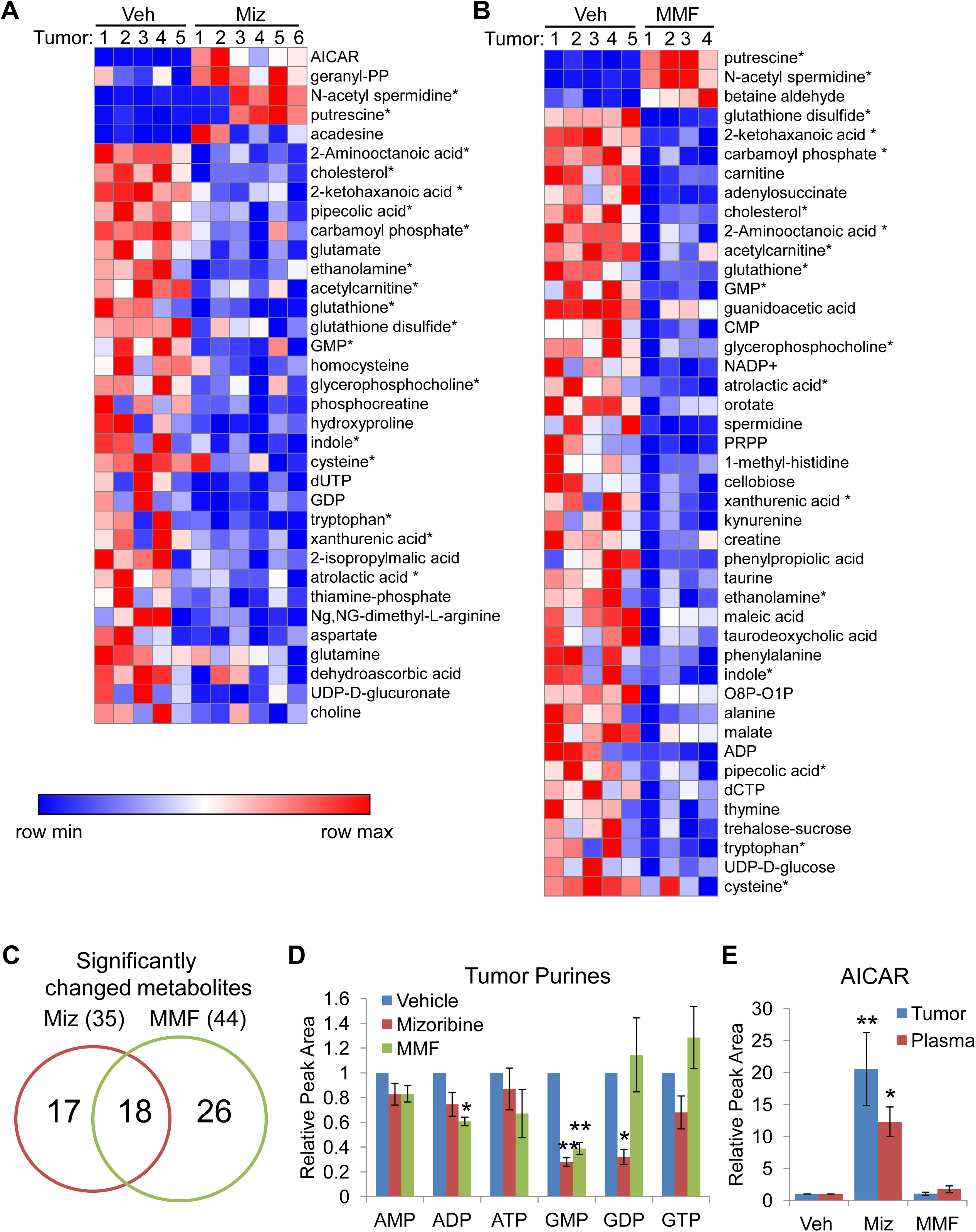
Effects of mizoribine and MMF on tumor metabolites. (A,B) Steady state LC-MS/MS-based metabolomics profiling on ELT3 xenograft tumors collected from the mice in Figure 3B on day 26, 3 hrs after the final treatment.(Vehicle: n = 5, Mizoribine: n = 6, MMF: n = 4). Row normalized heat maps are shown for all significantly changed metabolites (p < 0.05) in tumors from (A) mizoribine (Miz) or (B) MMF-treated mice compared to vehicle. Metabolites are grouped as increasing or decreasing relative to vehicle and then ranked from lowest p-value (top) to highest (bottom). Metabolites changed with both mizoribine and MMF are indicated with an asterisk. (C) Venn diagram showing the number of significantly changed metabolites in A,B. (D,E) Relative peak area values of (D) adenylate and guanylate nucleotides in tumors and (E) AICAR in tumors and plasma. Graphical data are presented as mean of indicated replicates ± SEM. *p < 0.05, **p < 0.005 by two-tailed Student’s t test.

In the 105K tumor model, 20 days of MMF treatment failed to reduce tumor GMP levels, but resulted in significant depletion of tumor GDP and GTP, as well as ADP and ATP (Supplementary Figure S3A). In these tumors, MMF caused only minor (1.6-fold) accumulation of AICAR (Supplementary Fig. S3B). As in the ELT3 tumor-bearing mice, mizoribine, but not MMF caused plasma AICAR accumulation (Supplementary Fig. S3C). Taken together, the decrease in tumor guanylates and increase in polyamines suggest that MMF treatment, like mizoribine, is directly affecting IMPDH in these two TSC tumor models. However, the greater depletion of tumor adenylates without effects on AICAR, with MMF treatment indicates distinct metabolic responses that might contribute to the differential anti-tumor responses measured between these two clinically used IMPDH inhibitors.

To more closely compare the plasma and tissue bioavailability of mizoribine and MMF, non-tumor bearing mice were analyzed after 7 days of treatment. Plasma concentrations of mizoribine and MPA were similar to those observed in our previous experiments prolonged (20 or 26 day) treatments (Fig. 2C, 3D, Supplementary Fig. S3D). Mizoribine, but not MMF treatment, caused AICAR accumulation to varying degrees in all tissues examined (plasma, liver, spleen, kidney, brain, lung, and heart), with the greatest accumulation observed in the liver, which is believed to be a major site of *de novo* nucleotide synthesis (Supplementary Fig. S4E)^47, 48^. Pharmacokinetic analysis after a single treatment with mizoribine or MMF revealed that despite a higher peak plasma concentration of MPA and slower clearance compared to mizoribine, plasma AICAR only accumulated after mizoribine treatment and did not return to baseline levels even after 24 or 48 hours (Supplementary Fig. S4F-I). Taken together, these data suggest that AICAR accumulation is a rapidly-induced, long-lived biomarker of IMPDH inhibition by mizoribine in plasma, normal tissues, and tumors.

### Mizoribine is superior to the maximum tolerated dose of MMF in an immunocompetent syngeneic TSC tumor model

Given the established immunosuppressive effects of IMPDH inhibitors, we compared mizoribine to MMF in an immunocompetent syngeneic xenograft tumor model, and also asked whether higher doses of MMF might have similar efficacy as mizoribine. We also controlled for the route of drug administration by delivering both mizoribine and MMF by oral gavage, recapitulating their most common delivery method in patients. We found that plasma mizoribine concentration and AICAR levels were higher when mizoribine was administered by oral gavage compared to the same dose administered via i.p. injection (Supplementary Fig. S4A,B), thus the dose was reduced for oral delivery to yield plasma concentrations similar to that obtained with i.p. injection (Fig. 5A). In a dosing pilot experiment, wild-type C57BL/6J mice were treated by oral gavage with vehicle, 50 mg/kg/day mizoribine, 100 mg/kg/day MMF, 300 mg/kg/day MMF, 500 mg/kg/day MMF, or 1 mg/kg Q2D rapamycin for 10 days. Although plasma MPA concentrations increased in a dose-dependent manner, the 5-fold increase in MMF dose yielded a less than 2-fold increase in plasma MPA concentration, suggesting a bioavailability limitation (Fig. 5A). Consistent with this, at lower doses, doubling the MMF dose from 25 to 50 mg/kg correspondingly doubled the resulting plasma MPA concentration, but doubling MMF from 50 to 100 mg/kg only increased plasma MPA levels by 1.4-fold (Supplementary Fig. S4C). Importantly, white blood cell counts indicated that the immunosuppressive effects of MMF were intact with this treatment and increased in a dose-dependent manner, with the immunosuppressive effects of rapamycin and mizoribine in this treatment regimen being comparable to 100 and 300 mg/kg MMF, respectively (Fig. 5B). Consistent with our findings in immunocompromised mice, mizoribine robustly increased plasma AICAR levels while MMF had only slight effects (Supplementary Fig. S4D).

**Figure 5.**
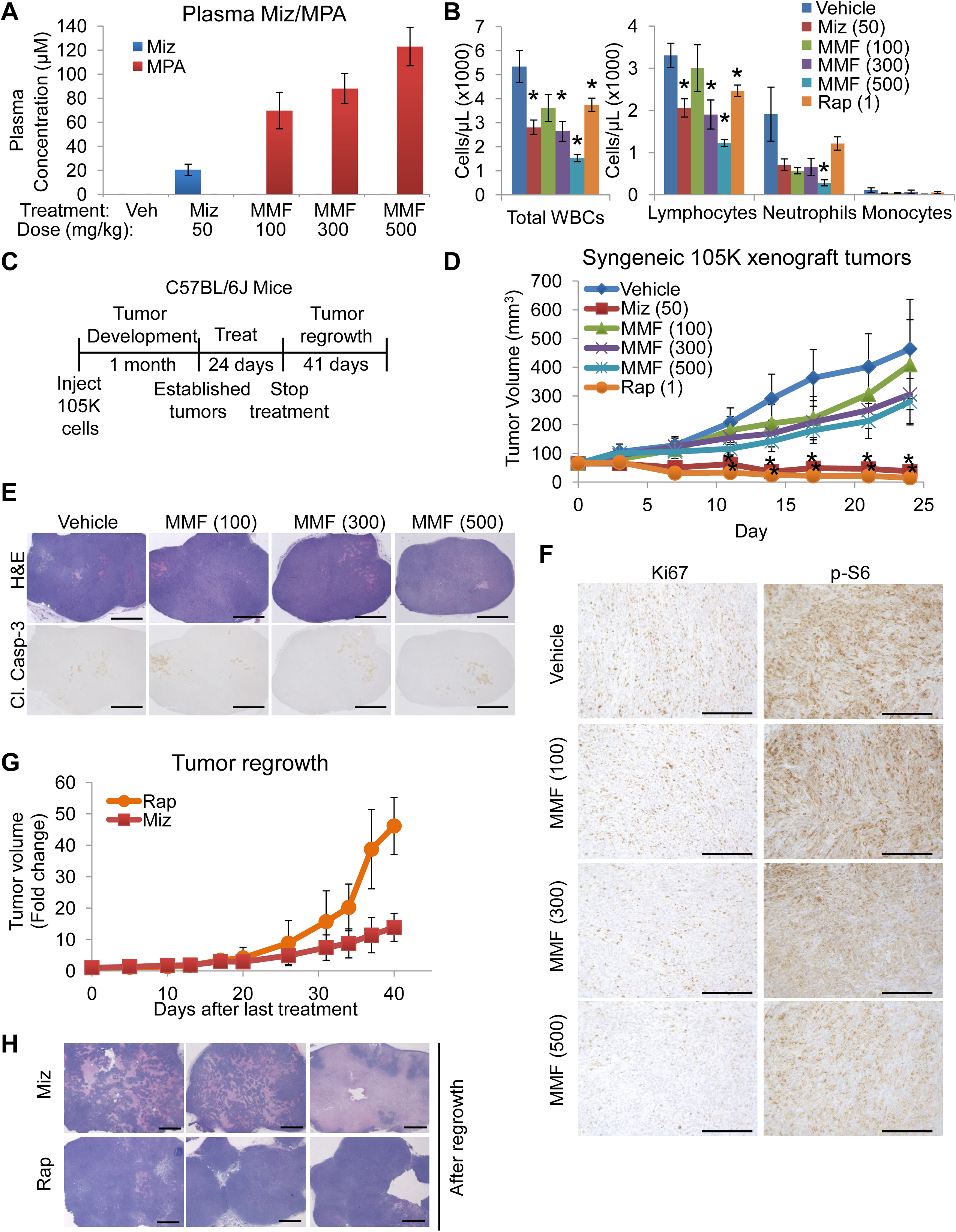
Mizoribine is superior to the maximum tolerated dose of MMF in an immunocompetent syngeneic xenograft model of TSC2-deficient tumor growth. (A,B) C57BL/6J mice were treated daily for 10 days with mizoribine (50 mg/kg by oral gavage) MMF (100, 300 or 500 mg/kg by oral gavage) or rapamycin (1 mg/kg/Q2D by i.p. injection). Blood and plasma were collected 2.5 hrs after the final treatment for measurement of (A) mizoribine and MPA concentrations in plasma by LC-MS/MS, and (B) white blood cell counts. n = 3 mice/group. (C) Experimental design used in (D-H). C57BL/6 mice bearing syngeneic 105K cell xenograft tumors were treated as in (A,B) for 24 days at which point mice were sacrificed or, for the mizoribine and rapamycin groups, treatments were discontinued and tumors were allowed to regrow. n = 6 mice/group. (D) Tumor volume during the treatment phase measured every 3^rd^ day. (E,F) H&E and immunohistochemistry staining of tumors resected 3 hrs after the final treatment (scale bars = 2 mm (E) and 0.25 mm (F)). (G) Fold change in tumor volume during the regrowth phase relative to the last day of treatment. (H) H&E staining of tumors resected after regrowth (scale bars = 2 mm). Graphical data are presented as mean of indicated replicates ± SEM. *p < 0.05, **p < 0.01 by two-tailed Student’s t test.

C57BL/6J mice bearing syngeneic 105K xenograft tumors were treated orally for 24 days with vehicle, mizoribine (50 mg/kg/day) or MMF (100, 300, or 500 mg/kg/day), compared to rapamycin (1 mg/kg MWF, i.p. injection), beginning when tumors first became palpable (Fig. 5C). Although MMF treatment slowed tumor growth in a dose dependent manner, the effect of mizoribine was far superior to even the highest MMF dose, shrinking tumors by approximately 40% over the 24-day treatment period (Fig. 5D). MMF treatment did not increase tumor apoptosis (Fig. 5E), but the modest decrease in tumor growth was accompanied by a dose-dependent decrease in the proliferation marker Ki67 and the mTORC1 signaling marker phospho-S6 (Fig. 5F). Mice treated with 500 mg/kg/day MMF experienced weight loss until treatment day 10 when every 3^rd^ day of treatment was skipped in this group to mitigate the toxicity, but weight loss was not observed in any other treatment group (Supplementary Fig. S4E). To directly compare the durability of the response to mizoribine to that of rapamycin, treatment was discontinued for these groups at day 24 and tumors were allowed to regrow (Fig. 5G). On the final day of treatment, the mean tumor volume in the mizoribine group was 2.4-fold larger than the rapamycin group. Therefore, although the absolute tumor volume between the mizoribine and rapamycin groups following the regrowth phase were similar (Supplementary Fig. S4F), tumors from mice previously treated with mizoribine regrew slower than those treated with rapamycin (Fig. 5G). Furthermore, consistent with our data from other tumor models (Fig. 2F), after up to 41 days of treatment withdrawal and tumor regrowth, the tumors from mice originally treated with mizoribine contained large areas of tumor necrosis and cell death, in stark contrast to tumors first treated with rapamycin (Fig. 5H).

We previously demonstrated that 1 month of mizoribine treatment reduces the volume and number of renal tumors that develop spontaneously in *Tsc2^+/-^* mice on the A/J strain background^18^. To determine whether MMF has anti-tumor efficacy in this genetic model, *Tsc2^+/-^* mice were treated for 1 month with MMF (75 mg/kg/day by oral gavage). No effect of MMF treatment on renal tumor volume was observed (Supplementary Fig. S4G). We were unable to compare MMF to mizoribine in these mice due to A/J strain-specific toxicity encountered when mizoribine was administered by oral gavage compared to our previous studies using i.p injection^18^, reflecting the observed increase in plasma mizoribine concentrations with oral delivery (Supplementary Fig. S4A,B). To determine whether this mizoribine toxicity is specific to *Tsc2^+/-^* mice, wild-type and *Tsc2^+/-^*mice on the A/J background and wild-type C57BL/6J mice were treated with mizoribine (30 mg/kg/day by oral gavage) for 1 week. Both the wild-type and *Tsc2^+/-^*A/J mice experienced similar weight loss, while the total body weight of C57BL/6J mice was unaffected, demonstrating that this effect is strain specific, not genotype specific (Supplementary Fig. S4H). Importantly, plasma mizoribine concentration was approximately 4-fold higher in the A/J mice, both wild-type and *Tsc2^+/-^*, compared to the C57BL/6J mice, despite receiving the same 30 mg/kg/day oral dose, thus providing a likely explanation for the increased toxicity in the A/J strain (Supplementary Fig. S4I).

### Adenosine kinase, required for mizoribine action, is expressed in human TSC-associated lesions

To inhibit IMPDH, the purine nucleoside analog mizoribine must be phosphorylated by adenosine kinase (ADK) to produce mizoribine-monophosphate (Fig. 6A)^18, 49^. siRNA-mediated knockdown of ADK in Tsc2-deficient 105K tumor cells reconstituted with either empty vector or wild-type TSC2 confirmed that ADK is required for the cell growth inhibitory effects of mizoribine, but not MPA (Fig. 6B,C), with remaining mizoribine activity reflecting incomplete knockdown of ADK (Fig. 6D). If mizoribine is to be repurposed for TSC-associated pathologies, ADK expression would be required in the target cells. IHC staining confirmed ADK expression localized to both the cytosol and nucleus (as reported previously^50^) in human TSC-associated pulmonary LAM and renal angiomyolipoma patient samples. ADK staining was concentrated in the distinct smooth muscle-like cells with spindle morphology that are characteristic of these lesions, which also stain strongly for S6 phosphorylation in adjacent serial sections (Fig. 6E,F). Thus, these lesions and other TSC-associated tumors may be candidates for mizoribine therapy.

**Figure 6.**
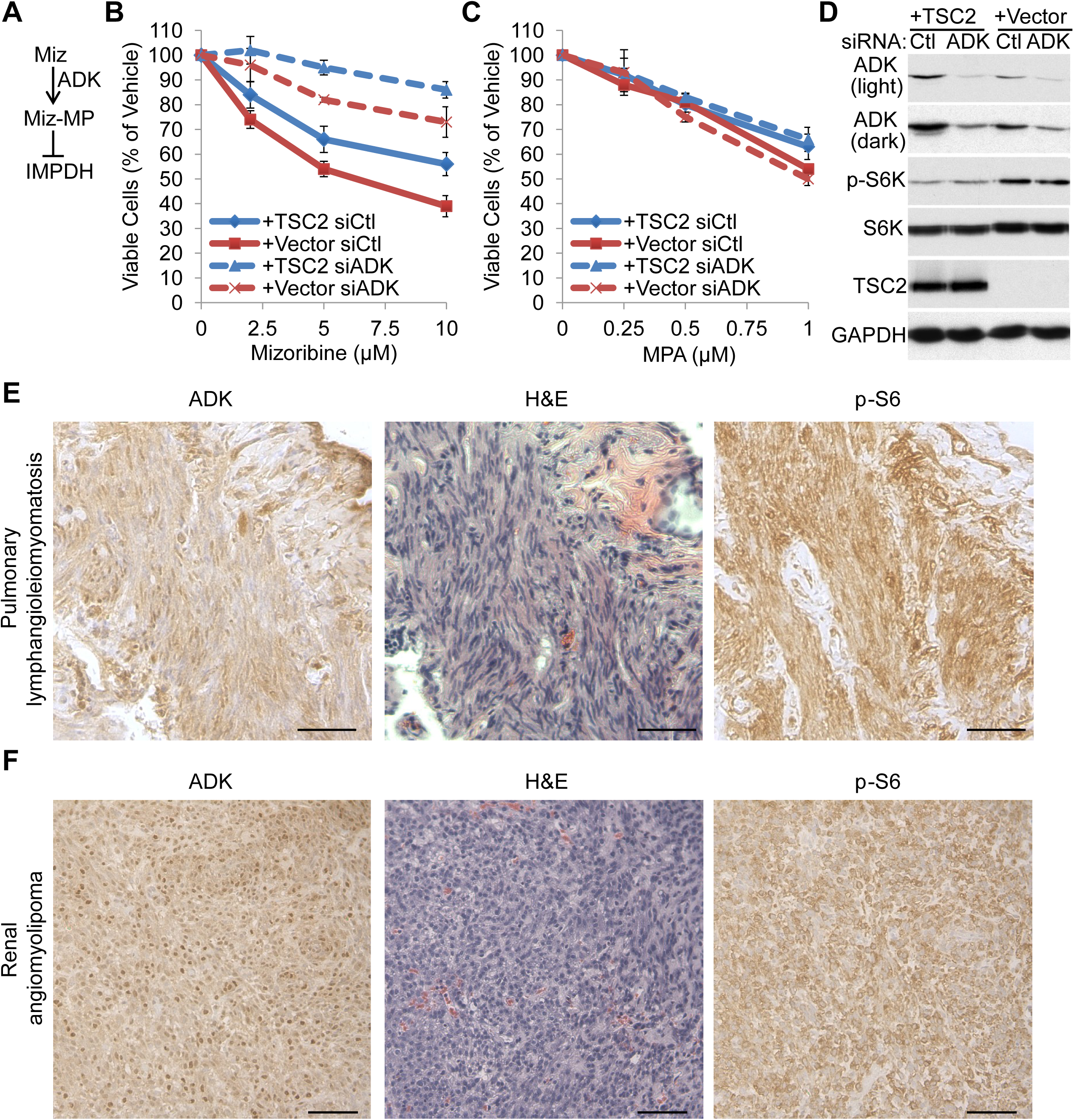
Adenosine kinase, required for mizoribine action, is expressed in human TSC-associated pulmonary lymphangioleiomyomatosis and renal angiomyolipoma. (A) Mizoribine is phosphorylated by adenosine kinase (ADK) to produce mizoribine monophosphate (Miz-MP) which inhibits IMPDH. (B-D) 105K cells stably reconstituted with empty vector or wild-type TSC2 were transfected with control (siCtl) or ADK-targeting siRNAs (siADK) and treated for 48 hrs with indicated concentrations of (B) mizoribine or (C) MPA. Viable cells were quantified by CellTiterGlo and graphed as percent of vehicle treated cells. n = 3 biological replicates. (D) Immunoblots showing ADK knockdown and TSC2 reconstitution efficiency. (E,F) H&E and immunohistochemistry staining in (E) human pulmonary lymphangioleiomatosis and (F) renal angiomyolipoma patient samples. Graphical data are presented as mean ± SEM. Scale bars = 100 µm

## Discussion

We previously found that uncontrolled mTORC1 activation increases cellular dependence on IMPDH for cell and tumor growth and viability by increasing the cellular demand for guanylate nucleotides required to support ribosomal RNA synthesis as part of mTORC1-driven ribosome biogenesis^18^. Here we directly compared clinically approved IMPDH inhibitors and find that mizoribine is a superior anti-tumor agent compared to mycophenolate in TSC2-deficient tumors, which are driven by robust mTORC1 activation. Collectively, we observe strong anti-tumor effects of mizoribine compared to mycophenolate in four TSC2-deficient tumor models: 105K and ELT3 xenograft tumors in immunocompromised mice (Figure 2,3), syngeneic 105K xenograft tumors in immunocompetent mice (Figure 5), and sporadic renal tumors in *Tsc2^+/-^* mice (Supplementary Fig. S4G and Ref^18^). These preclinical studies suggest mizoribine could be an effective treatment in tumors with active mTORC1, including those in TSC patients. Mizoribine has a potential advantage over rapamycin and its analogs, which are now often used for treating tumors in TSC patients, in its ability to induce tumor cell death and correspondingly slow the rate of tumor regrowth when treatment is discontinued. Importantly, decades of clinical mizoribine use in Asia indicate that it has a safety profile in humans that is similar or favorable to mycophenolate and rapamycin^23, 51–53^.

It is interesting to note that tumors lacking ADK expression could potentially be an exception to the findings presented here, in which case MMF might have an advantage over mizoribine due to the inability of those cells to convert mizoribine to the active mizoribine-monophosphate. Thus, we confirmed ADK expression in human TSC-associated lymphangioleiomyomatosis and renal angiomyolipoma samples. We also identified accumulation of the purine synthesis intermediate AICAR as a metabolic biomarker of mizoribine efficacy in cells, tumors, normal tissues and plasma, which could be used to monitor the activity of mizoribine.

Although mizoribine and mycophenolic acid both inhibit IMPDH and are used for similar clinical indications, we find that they have differential effects on signaling and metabolism *in vivo*. Unlike mizoribine, and in contrast to MPA in cultured cells, *in vivo* MMF treatment causes depletion of adenylate nucleotides and does not cause accumulation of AICAR. The mechanistic nature of these differential effects and whether they limit the specificity and efficacy of MMF in TSC tumors is currently unknown. Another difference between mizoribine and MMF activity that could influence its anti-tumor efficacy is that MMF was found to partially inhibit mTORC1 signaling. Our previous studies demonstrated that sensitivity to IMPDH inhibitors requires robust and sustained mTORC1 activation, with consequent elevated rates of rRNA synthesis^18^.

It is remarkable that available inhibitors of other essential enzymes in the *de novo* or salvage nucleotide synthesis pathways, including several compounds in clinical use, failed to show the selective inhibitory effects on growth and viability of TSC2-deficient cells that we observe with IMPDH inhibitors. These results suggest that targeting guanylate synthesis, specifically, offers a unique ability to exploit the increased demand for nucleotides that comes with mTORC1-driven ribosomal RNA synthesis. Guanylate nucleotides are unusually enriched in ribosomal RNA, accounting for 34% of nucleotides in the mature mammalian ribosome and 37% of nucleotides in the 45S pre-rRNA that is processed into the mature 18S, 5.8S, and 28S rRNAs. Correspondingly, the rate of incorporation of available GTP into rRNA is greater than ATP, UTP or CTP^20^. Free intracellular guanylates are maintained at substantially lower concentrations than adenylates, and are also lower than pyrimidines in many cell types ^54–58^. Thus, lower basal levels combined with greater enrichment in rRNA could underlie the distinct importance of guanylate synthesis in cells with elevated rates of rRNA synthesis, and provide a mechanism for the selective vulnerability of cells with elevated mTORC1 signaling to IMPDH inhibitors, relative to those inhibiting the synthesis of other nucleotides.

Like mTORC1, the oncogene Myc also drives both ribosome biogenesis and *de novo* nucleotide synthesis^59, 60^. Dependence on IMPDH correlates with Myc expression in small cell lung cancer, where mizoribine demonstrated anti-tumor efficacy in preclincal tumor models^19^. IMPDH dependence correlating with a high rate of rRNA synthesis was also recently demonstrated in glioblastoma where xenograft tumor growth was slowed by MMF and completely blocked by genetic knockout of IMPDH1 and IMPDH2 ^20^. MMF has also demonstrated efficacy in xenograft tumors established using other human cancer cell lines^21, 22^. This growing body of evidence suggests that IMPDH inhibitors could potentially be repurposed as anti-tumor agents, and our study indicates that mizoribine should be considered a leading candidate amongst these drugs.

## Acknowledgements

We thank David J. Kwiatkowski, Gerta Hoxhaj, Issam Ben-Sahra, Hilaire Lam and Min Yuan for helpful discussions and technical assistance. We also thank the Tuberous Sclerosis Alliance TSC Preclinical Consortium for experiments in *Tsc2^+/-^* mice.

