## Supplemental Figures for "IMPDH inhibitors for anti-tumor therapy in tuberous sclerosis complex"

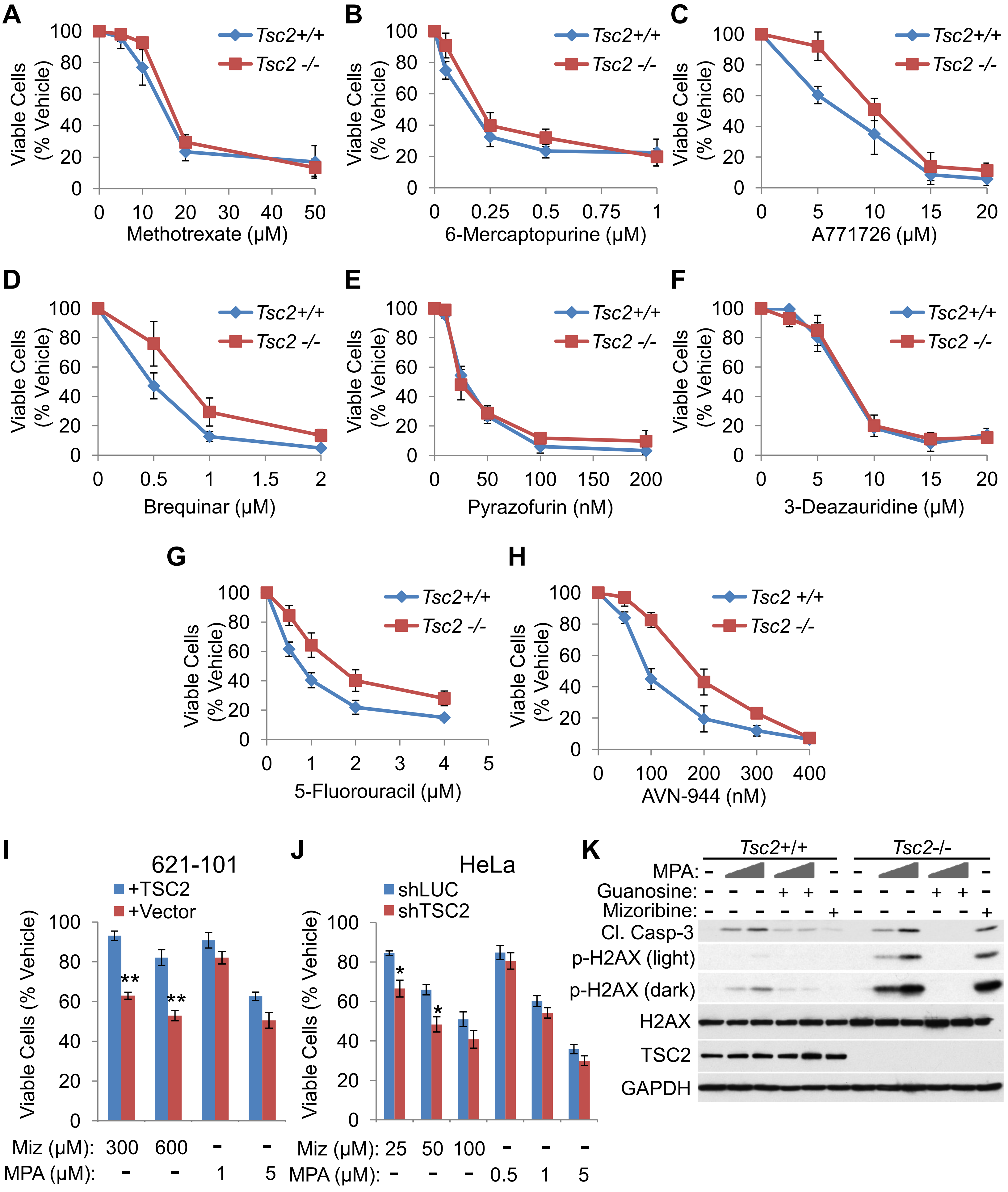


**Supplemental Figure 1**

**Figure S1. Supplementary data supporting Figure 1**

(A-H) Viable cell counts in littermate derived *Tsc2*$^{+/+}$ and *Tsc2*$^{-/-}$ MEFs treated with vehicle or indicated concentrations of indicated compounds for 72 hrs and graphed as percent of vehicle treated cells. n = 3 biological replicates. (I,J) Viable cell counts in (I) *TSC2*$^{-/-}$ 621-101 human renal angiomyolipoma-derived cells stably reconstituted with empty vector or wild-type TSC2, or (J) HeLa cells with stable shRNA-mediated knock down of TSC2 or control (luciferase) treated for 48 hrs with indicated concentrations of mizoribine or MPA and graphed as percent of vehicle treated cells. n = 3 biological replicates. (K) Immunoblots on *Tsc2*$^{+/+}$ and *Tsc2*$^{-/-}$ MEFs treated for 48 hrs as indicated with MPA (200 or 300 nM), guanosine (guan, 50 μM) or mizoribine (miz, 2 μM).

Graphical data are represented as mean of indicated replicates, error bars represent ± SEM. *p < 0.05, **p < 0.01 by two-tailed Student's t test.

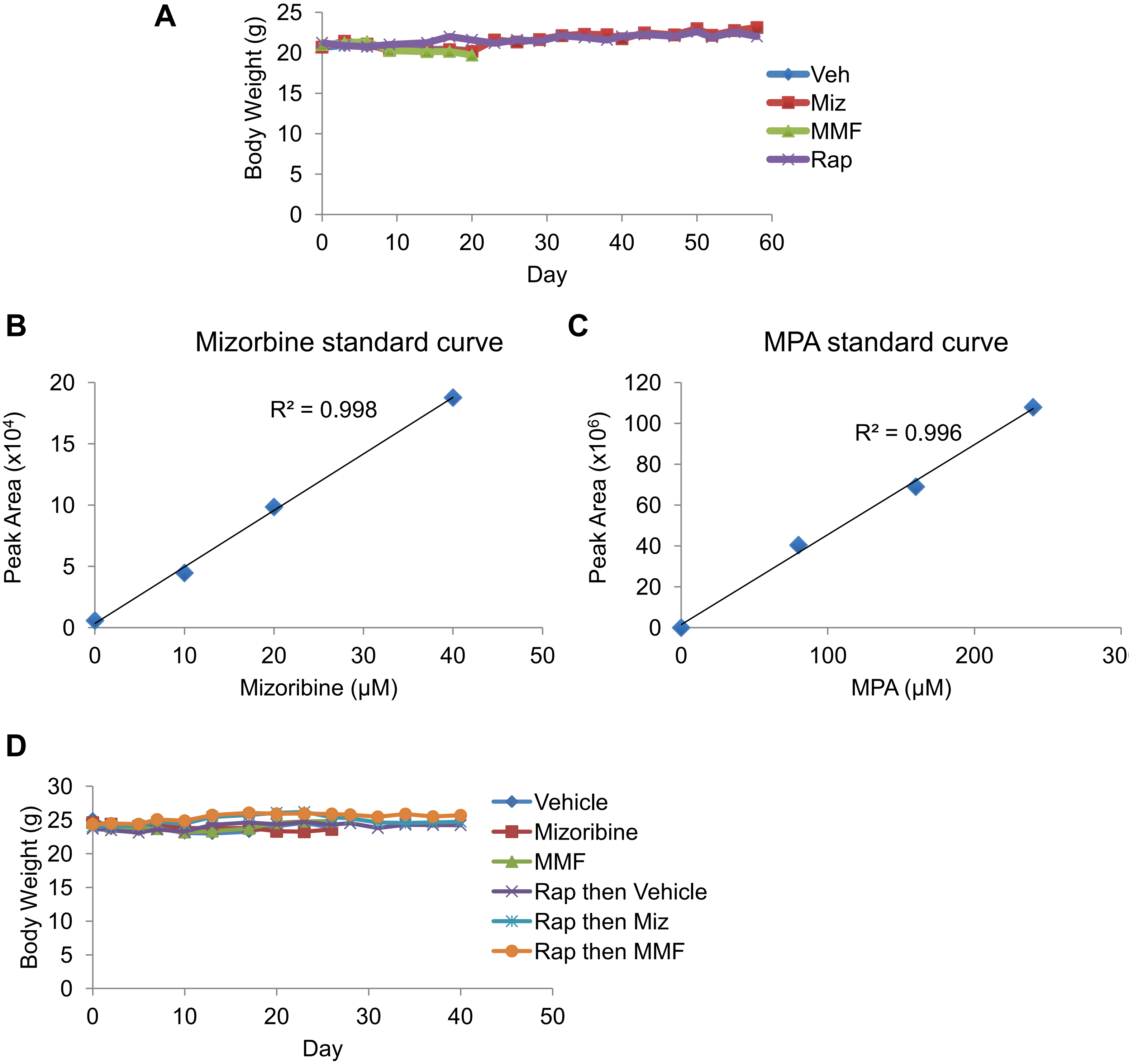


**Supplemental Figure 2**

**Figure S2. Supplementary data supporting Figures 2,3**

(A) Total body weight of mice in Figure 2B. (B,C) Mizoribine and MPA standard curves used to quantify plasma concentrations in Figure 2C. Mizoribine or MPA were added into plasma from untreated mice at indicated final concentrations. Metabolites were extracted from these standards alongside plasma from vehicle, mizoribine and MMF treated mice, and mizoribine and MPA were measured by LC-MS/MS. (D) Total body weight of mice in Figure 3B.

Graphical data are represented as mean of indicated replicates, error bars represent ± SEM.

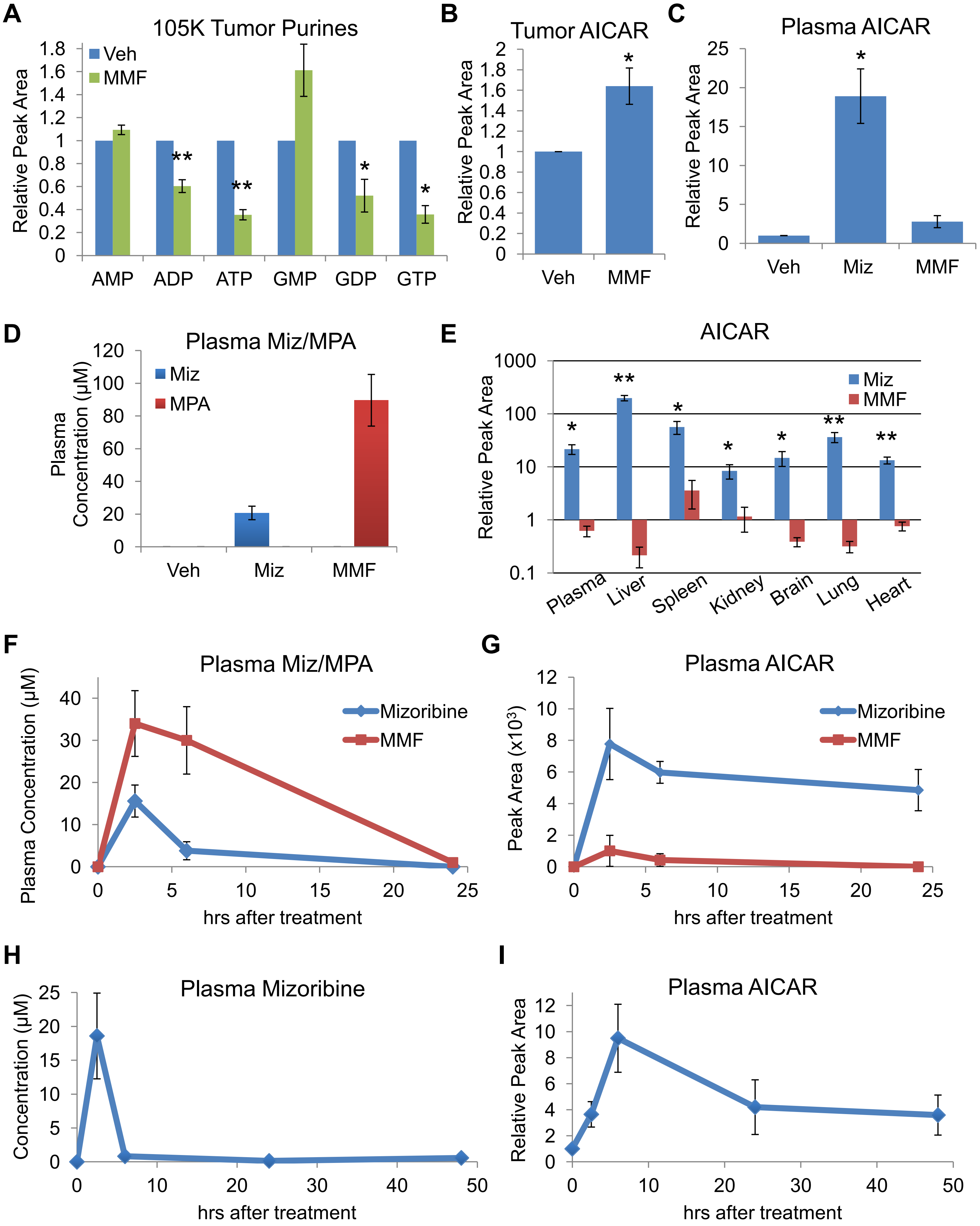


**Supplemental Figure 3**

**Figure S3. Supplementary data supporting Figure 4**

(A-C) Steady state LC-MS/MS-based metabolomics profiling on 105K xenograft tumors or plasma collected from the mice in Figure 2B. (Vehicle: n = 6, MMF n = 6). Relative peak area values of (A) tumor adenylates and guanylates, (B) tumor AICAR, or (C) plasma AICAR. (D,E) Wild type C57BL/6J mice were treated daily by oral gavage with vehicle, mizoribine (45 mg/kg) or an equimolar dose of MMF (75 mg/kg) for 7 days. n = 4 mice/group. (D) Mizoribine and MPA concentrations in plasma collected 2.5 hrs after the final treatment. (E) AICAR levels measured by LC-MS/MS in the indicated tissues with peak area values graphed on a log10 scale relative to vehicle. (F,G) NSG mice were given a single treatment of mizoribine (50 mg/kg) or MMF (50 mg/kg) by oral gavage and plasma was collected at indicated time points for LC-MS/MS measurement of (F) mizoribine and MPA concentration and (G) AICAR levels. n = 3 mice for each time point. (H,I) Wild type C57BL/6J mice were given a single treatment of mizoribine (100 mg/kg by i.p. injection) and plasma was collected at indicated time points for LC-MS/MS measurement of (H) mizoribine concentration and (I) AICAR levels. n = 3 mice for each time point.

Graphical data are represented as mean of indicated replicates, error bars represent ± SEM. *p < 0.05, **p < 0.01 by two-tailed Student's t test.

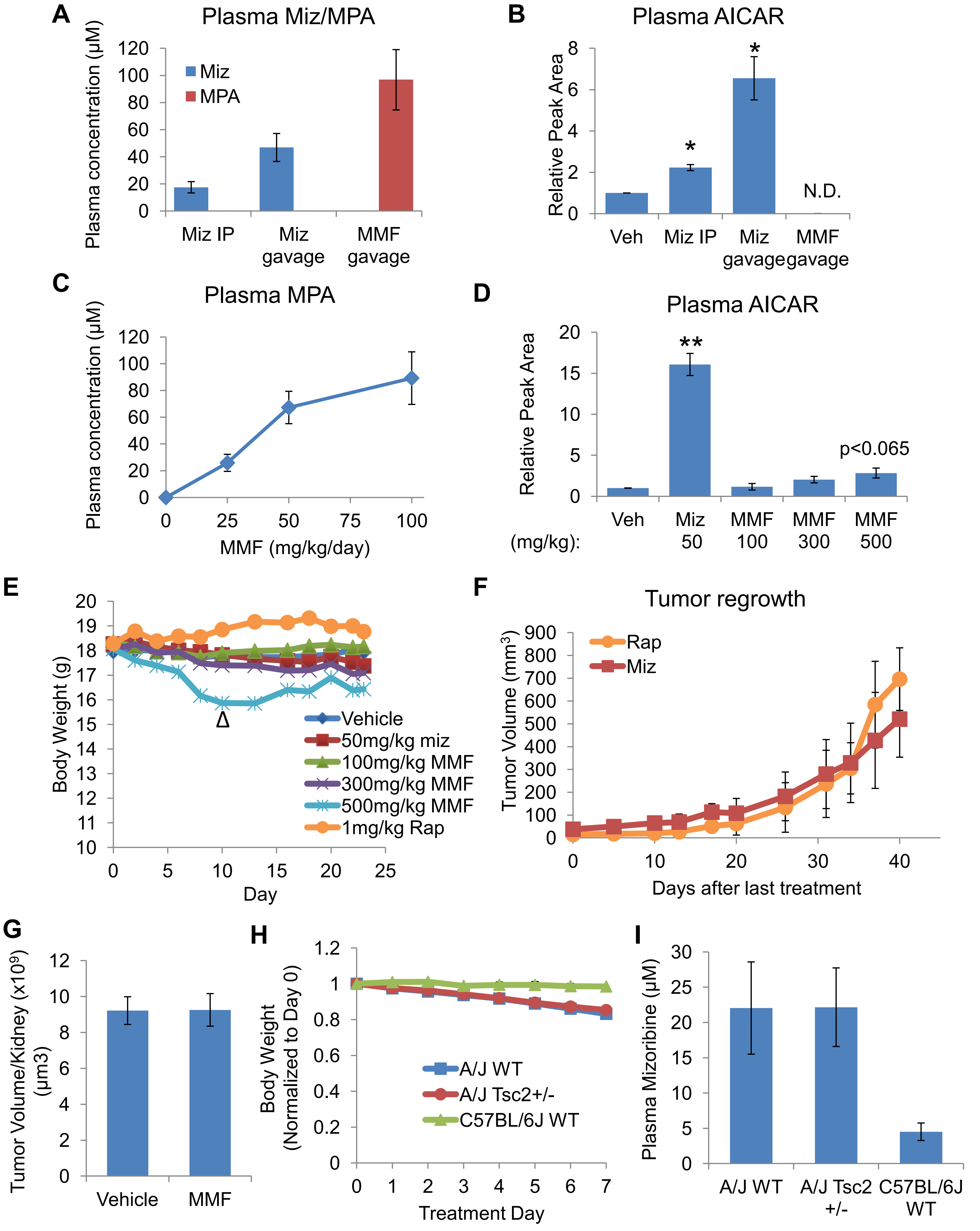


**Supplemental Figure 4**

**Figure S4. Supplementary data supporting Figure 5**

(A,B) Wild type C57BL/6J mice were treated daily with mizoribine (100 mg/kg) by i.p. injection, mizoribine (100 mg/kg) by oral gavage or an equimolar dose of MMF (167 mg/kg) by oral gavage for 3 days. (A) Mizoribine and MPA concentration and (B) AICAR levels were measured by LC-MS/MS in plasma collected 2.5 hrs after the final treatment. n = 3 mice/group. N.D. = not detected. (C) MPA concentration in plasma from C57BL/6J mice treated daily by oral gavage with indicated doses of MMF for 4 days. n = 3 mice/group. (D) Relative AICAR levels in plasma from Figure 5A, measured by LC-MS/MS. (E) Total body weight of mice in Figure 5D. Every 3$^{rd}$ day of treatment was skipped in the 500 mg/kg MMF group beginning on day 10 (indicated by Δ) to mitigate weight loss. (F) Volume of tumors measured every 3$^{rd}$ day during the regrowth phase in Figure 5F. (G) *Tsc2*$^{+/-}$ mice on the A/J strain background were treated with vehicle or MMF (75 mg/kg/day) by oral gavage for 1 month beginning at 7 months of age. n = 8 mice/group (H,I) Mice of the indicate genotypes and strain backgrounds were treated with mizoribine (30 mg/kg/day) for 7 days by oral gavage. n = 4 mice/group. (H) Total body weight measured daily. (I) Mizoribine concentration in plasma collected 2.5 hrs after the final treatment. Graphical data are represented as mean of indicated replicates, error bars represent ± SEM. *p < 0.05 by two-tailed Student's t test.
